# FIGARO: An efficient and objective tool for optimizing microbiome rRNA gene trimming parameters

**DOI:** 10.1101/610394

**Authors:** Michael M. Weinstein, Aishani Prem, Mingda Jin, Shuiquan Tang, Jeffrey M. Bhasin

## Abstract

**Summary:** Microbiome studies continue to provide tremendous insight into the importance of microorganism populations to the macroscopic world. High-throughput DNA sequencing technology (i.e., Next-generation Sequencing) has enabled the cost-effective, rapid assessment of microbial populations when combined with bioinformatic tools capable of identifying microbial taxa and calculating the diversity and composition of biological and environmental samples. Ribosomal RNA gene sequencing, where 16S and 18S rRNA gene sequences are used to identify prokaryotic and eukaryotic species, respectively, is one of the most widely-used techniques currently employed in microbiome analysis. Prior to bioinformatic analysis of these sequences, trimming parameters must be set so that post-trimming sequence information is maximized while expected errors in the sequences themselves are minimized. In this application note, we present FIGARO: a Python–based application designed to maximize read retention after trimming and filtering for quality. FIGARO was designed specifically to increase reproducibility and minimize trial-and-error in trimming parameter selection for a DADA2–based pipeline and will likely be useful for optimizing trimming parameters and minimizing sequence errors in other pipelines as well where paired-end overlap is required.

**Availability and implementation:** The FIGARO application is freely available as source code at https://github.com/Zymo-Research/figaro.

## 1. INTRODUCTION

The study of microbial communities has proven critical to further understanding of a wide variety of topics ranging from agriculture to tumor immunology^1,2^. Targeted sequencing, particularly of the ribosomal RNA (rRNA) gene sequence, remains a popular and efficient option for identifying different microbes present and assessing the microbial composition of a sample^3^. Illumina provides the industry leading platform for high-throughput sequence analysis with an error model that is well-studied^4^. Due to the expected presence of errors generated from Illumina sequencing, different methods have been developed to compensate for these errors in an effort to minimize spurious identification of novel microbial community members arising from sequencing artifacts. Initially, clustering based upon a fixed threshold (generally 3%) of dissimilarity (often called the OTU approach) was the standard^5^. These methods improved specificity in microbiome analyses at the cost of resolution. Recently, there has been a demand for higher resolution targeted sequencing approaches for species- and strain-level identification of microbes. Fulfilling this demand requires statistical methods to correct sequencing errors, with DADA2^6^ and Deblur^7^ being common solutions. When utilizing either of these methods for paired-end reads, it is necessary to merge the read pairs prior to analysis; this is only possible if sufficient length remains on both reads after trimming to cover the entire amplicon with sufficient overlap length. FIGARO is a Dockerized Python-based application that can select optimal trimming sites for paired-end data that allows for minimizing expected sequencing errors while maximizing read retention and maintaining sufficient read length for downstream merging.

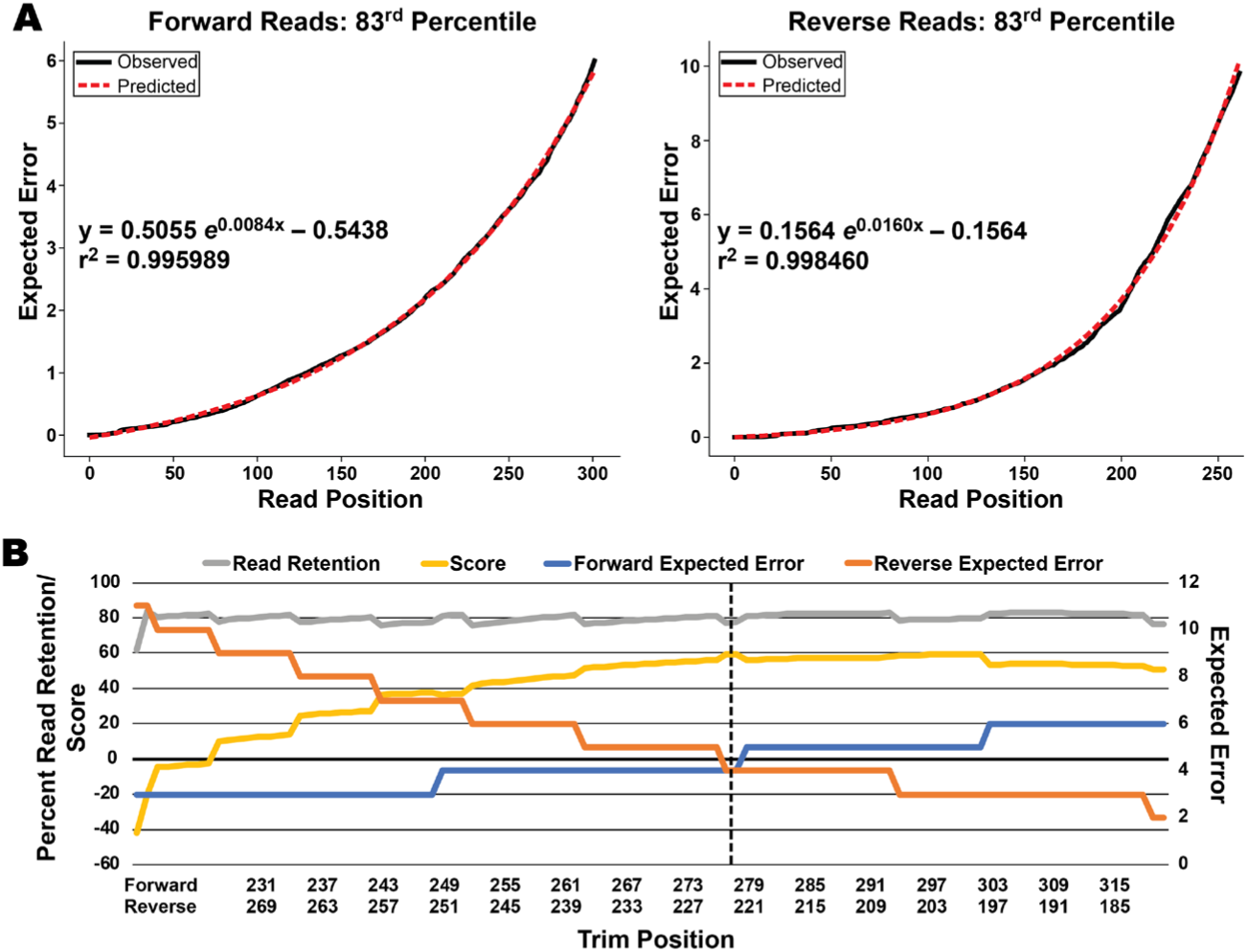
Panel A: Fitting an exponential regression to the 83^rd^ percentile for cumulative expected error values across multiple samples from a single sequencing experiment on a MiSeq. The high (>0.99) r^2^ value in both directions is representative of what was often observed with this model. Panel B: A plot showing the percent read retention, trimming site scores, and forward and reverse expected error allowances for a set of 16S rRNA gene sequences covering the V3 and V4 regions generated on a MiSeq. The vertical dashed line represents the trimming site recommended by FIGARO, providing minimal expected error allowances in both directions while still preserving the expected percentage of reads. Panel B: A plot showing the percent read retention, trimming site scores, and forward and reverse expected error allowances for a set of 16S rRNA gene sequences covering the V3 and V4 regions generated on a MiSeq. The vertical dashed line represents the trimming site recommended by FIGARO, providing minimal expected error allowances in both directions while still preserving the expected percentage of reads.

## 2. DESCRIPTION AND USAGE OF FIGARO

### User inputs

The user must provide the total length of their amplicon as well as a folder containing untrimmed paired-end read data in FASTQ format. Optional inputs include specifying the name of the output file, specifying input and output directories, subsampling rate for the FASTQ files, minimum overlap between the paired-end reads, and the expected error percentile to use in filtering at each tested position.

### Modeling expected errors by read position

Like DADA2^6^, Figaro utilizes the cumulative expected errors for quality filtering^8^. Because the rate of expected error accumulation across reads can vary between devices, runs on the same device, and read directions on the same run, an exponential regression model is built for each set of reads analyzed using the generic equation *y* = *a* · *e*^*bx*^ +*c.* This model describes the *n*-th percentile for expected errors at a given position in the read (with a default value of *n*=84, corresponding to 1 standard deviation above the mean) and is found using the *curve_fit* function in *Scipy*’s *optimize* package^9^. This regression model was initially selected based upon observed data strongly suggesting an exponential model and reinforced by subsequent applications of this model often yielding R^2^-values greater than 0.99 for both read directions (Figure panel A).

### Selecting candidate trimming parameters

Given an expected amplicon size and minimum overlap between the forward and reverse reads, calculating the required combined read-length for both reads can be done by summing the amplicon size and overlap length. The candidate trimming sites are found by determining all combinations of forward and reverse trimming positions that will yield the required combined read-length. Maximum expected error values are calculated for forward and reverse reads based upon the candidate trimming position and the previously determined exponential regression model.

### Calculating read retention post-filtering

To predict the effect of filtering on read retention, the *Numpy* library is used to construct an array of cumulative expected error values for all potential trimming positions within every read in the supplied data set^10^. Subsampling can be used to improve speed and decrease the memory footprint of FIGARO while still achieving accurate results, with the default subsampling behavior being determined by the size of the input data. The forward and reverse expected error arrays are then iterated over at the candidate trimming sites to quantify how many reads would be expected to remain following quality filtering.

### Scoring candidate trimming sites

Candidate trimming sites are scored based upon their rate of read retention with a penalty subtracted for expected errors greater than one as follows, where RR is the percent read retention and EE represents the expected error:

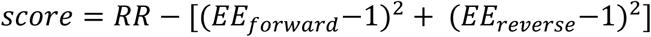

### Outputs

The primary output of FIGARO is a list of potential trimming site parameters sorted from highest-scoring to lowest. These results can be output to both the console for manual review, or to a JSON-formatted text file for parsing if using an automated pipeline. Additionally, FIGARO will generate figures describing the expected error regression models for both forward and reverse reads.

## 3. RESULT

Utilizing FIGARO, combinations of trimming sites were found that could be used while keeping expected error allowances in each paired read to a minimum and still allowing the expected percentage of reads to pass filtering (Figure panel B). Utilizing FIGARO on reads generated using the ZymoBIOMICS™ Microbial Community Standard—a mock microbial community with known organisms present in a known proportion—allowed us to automatically select trimming parameters in under a minute on a laptop computer. These parameters, utilized in a typical DADA2 pipeline, were capable of detecting all expected organisms in appropriate proportions. Analyzing untrimmed reads in the same pipeline only detected a single organism and determining appropriate trimming parameters with an experienced technician required several minutes of observation and multiple iterations of trial-and-error.

## 4. CONCLUSION

Sequence trimming is a critical step in a high-resolution microbiome analysis pipeline that is sometimes left to trial and error or the best guess of the pipeline operator. While this is likely adequate for shorter amplicons (such as those covering a single variable region of the 16S rRNA), longer amplicons—often covering multiple variable regions of the 16S rRNA gene—are often approaching the technical capabilities of the sequencing platform and require more intensive optimization. FIGARO models the error rate for each sequencing run in each direction to find optimal trimming sites that will maximize read retention after filtering while removing some lowest-quality percentile of reads.

